# The mechanics of predator-prey interactions: first principles of physics predict predator-prey size ratios

**DOI:** 10.1101/313239

**Authors:** Sébastien M. J. Portalier, Gregor F. Fussmann, Michel Loreau, Mehdi Cherif

**Affiliations:** Department of Biology, McGill University, Montreal, QC, Canada; Centre for Biodiversity Theory and Modelling, Theoretical and Experimental Ecology Station, CNRS and Paul Sabatier University, 09200 Moulis, France; Department of Ecology and Environmental Sciences, Umeå University, Sweden

**Keywords:** predation, trophic link, body size, energy, mechanics

## Abstract

1. Robust predictions of predator-prey interactions are fundamental for the understanding of food webs, their structure, dynamics, resistance to species loss and invasions and role in ecosystem functioning. Most current food web models are empirically based. Thus, they are sensitive to the quality of the data, and ineffective in predicting non-described and disturbed food webs. There is a need for mechanistic models that predict the occurrence of a predator-prey interaction based on the traits of organisms and the properties of their environment.
2. Here, we present such a model that focuses on the predation act itself. We built a Newtonian, mechanical model for the processes of searching, capture and handling of a prey item by a predator. Associated with general metabolic laws, we predict the net energy gain from predation for pairs of predator and prey species depending on their body sizes.
3. Predicted interactions match well with data from the most extensive predator-prey database, and overall model accuracy is greater than the niche model.
4. Our model shows that it is possible to accurately predict the structure of food webs using only a few ecomechanical traits. It underlines the importance of physical constraints in structuring food webs.

## 1 Introduction

Predicting predator-prey interactions accurately is fundamental. The dynamics of food webs depend critically on their structures (Allesina & Tang, 2012). Moreover, the fate of native and invasive species depends on the network of interactions in which they are embedded (Romanuk et al., 2009). There is also increased awareness that ecosystem functioning itself depends on the structure of food webs (Thompson et al., 2012). It is thus important to understand what determines the occurrence of pairwise predator-prey interactions and by extension, the structure of food webs.

Most historical and contemporary models that predict the structure of food webs are empirically based. They derive rules from the regularities observed in well-studied food webs; devise statistical models that can reproduce these regularities in simulated food webs; and test the capacity of these statistical models to predict the structure of newly described food webs (Cohen & Newman, 1985; Williams & Martinez, 2000; Eklöf et al., 2013; Gravel, Poisot, Albouy, Velez, & Mouillot, 2013). While these models often succeed in faithfully replicating the patterns from which they are constructed, their performance worsens when it comes to other features of food webs (Williams & Martinez, 2008). Moreover, there are still limits to how accurate and detailed one can go in the description of food webs, despite steady improvements in the quality and quantity of food web data (Evans, Kitson, Lunt, Straw, & Pocock, 2016). As a result, most food web data are still irremediably spatially, temporally and/or taxonomically aggregated (Martinez, Hawkins, Dawah, & Feifarek, 1999). Hence, statistical modelling approaches describe reasonably well food webs similar to those on which they have been built and trained, but they might have issues to describe other food webs, knowing that discrepancies exist among ecosystems (Yvon-Durocher et al., 2011).

Thus, a complementary, mechanistic approach to food web modelling is needed, one in which pairwise interactions can be predicted from species traits and properties of the environment, and the food web emerges from combining all the potential pairwise interactions between the species present (Stouffer, 2010).

One trait that has focused the attention of food web ecologists, and for good reasons, is body size (Cohen, Pimm, Yodzis, & Saldaña, 1993; Loeuille & Loreau, 2005; Petchey, Beckerman, Riede, & Warren, 2008; Gravel, Poisot, Albouy, Velez, & Mouillot, 2013). These studies have made great strides to reveal the role of size in structuring food webs, including its role in determining functional responses and interactions strengths (Brose, 2010). But here again, the patterns of prey-to-predator body size ratios and allometries used are empirical, and thus they do not offer any mechanistic underpinning. Hence, the question of the factors that determine the size of the prey selected by a predator of a given body size remains incompletely understood as well as the mechanisms by which these factors operate.

To answer this question, we decided to adopt a mechanistic approach, concentrating on the core of predator-prey interactions, the act of predation itself, represented by the local search, capture and handling of one prey item by one given predator. The originality of our approach is to consider that predation is by essence a mechanical interaction (Figure 1): the predator must set itself in motion to search and capture the prey, while the prey moves to avoid capture. The act of handling involves mechanical motion as well since the predator must maintain its position in the water or air column while eating its prey. We used Newton’s laws of mechanics associated with optimization techniques as a basis to estimate encounter rates, capture probabilities and handling times for all predator-prey pairs within a realistic range of body sizes. Combined with general laws about metabolic expenses in organisms, we then used this mechanical model to calculate an energy budget for the predator during predation, and thus determine prey profitability. One advantage of our model is that only the body sizes of the species are used as inputs. No other parameter is fit to the data.

**Figure 1:**
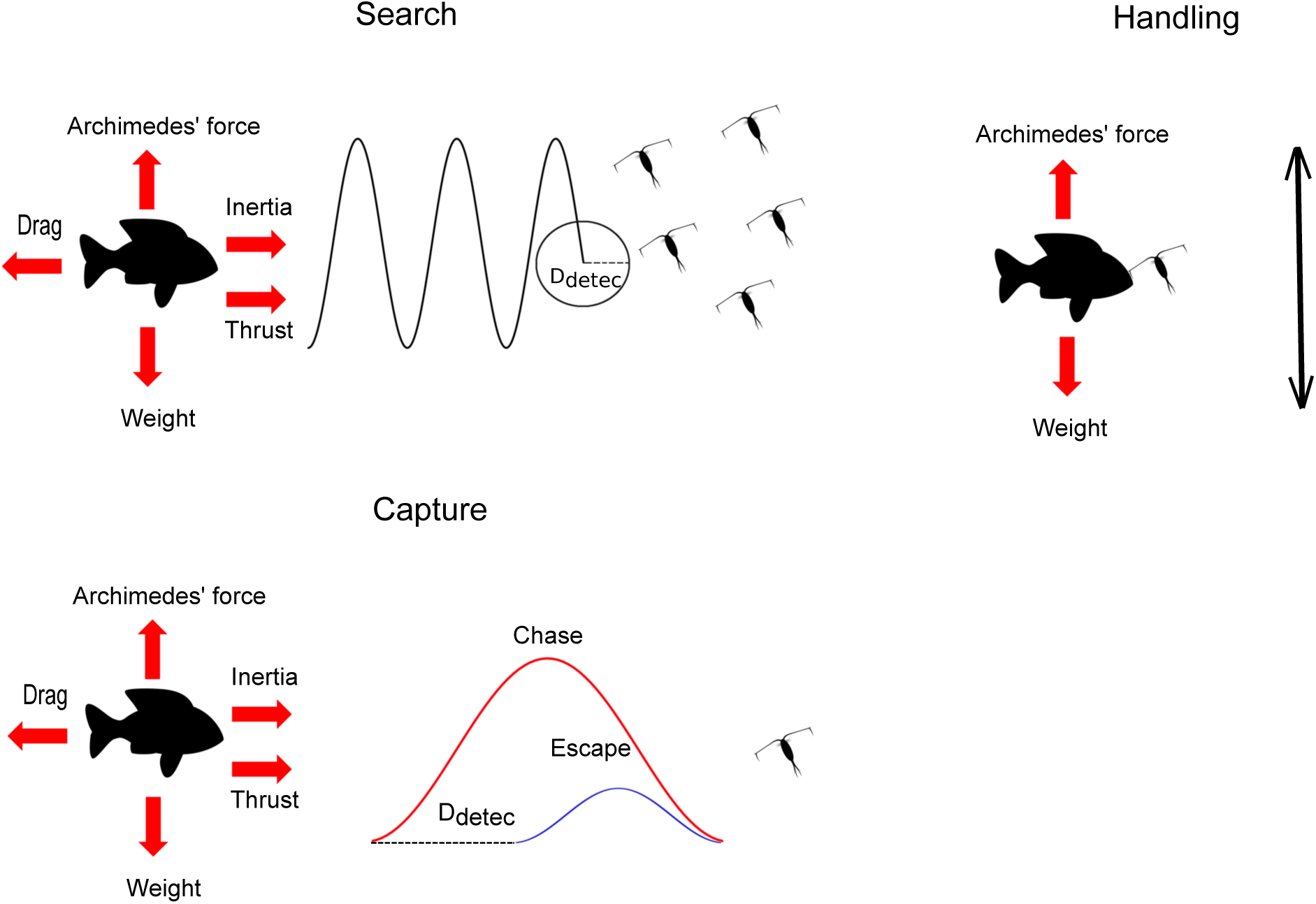
Main features of the model. The predator needs to spend energy to hover despite its weight (due to gravity), but it benefits from Archimedes’ force. Predation is split into three sequences. First, the predator searches prey. Motion implies interplay between mechanical thrust, inertia and drag. Encounter is constrained by predator’s detection distance (*D_detec_*), and prey abundance. A successful encounter leads to the capture sequence: the predator moves to seize the prey, while the prey tries to escape. In case of a successful capture, the predator needs to handle the prey during handling time (consumption and digestion): the predator needs to maintain hovering (lifting itself and the prey).

Thanks to the mechanical underpinning of our model, we can predict prey body sizes and profitability for both pelagic and flying predators. Including mechanics in ecological studies allows unifying approaches and comparisons among systems rather than being restricted to a specific habitat (Webb, 2012). Hence, our model opens the door to a bottom-up prediction of the structure of food webs in diverse physical habitats, based only on a few mechanical traits of both predators and their prey.

## 2 Methods

The model calculates a net energetic gain (*G*) for each predator-prey interaction:

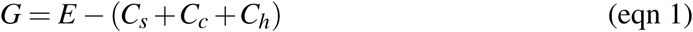

where *E* is energy received from the prey, *C_s_*, *C_c_* and *C_h_* are the costs for searching, capturing and handling the prey respectively. Most of parameters used in the model scale with body size. *M_pred_* refers to predator mass, while *M_prey_* refers to prey mass. The model is static and only includes energy allocations related to predation (i.e, no predator growth or reproduction).

### 2.1 Physical parameters

The model considers two different media (air and water). Parameters used by the model are acceleration due to gravity (*g*), body density (*ρ_b_*), medium density (*ρ_m_*), and medium dynamic viscosity (*μ*, see Table 1). Motion involves calculation of the drag coefficient in order to estimate speeds and related power outputs (see Supplementary Methods 1).

**Table 1:**
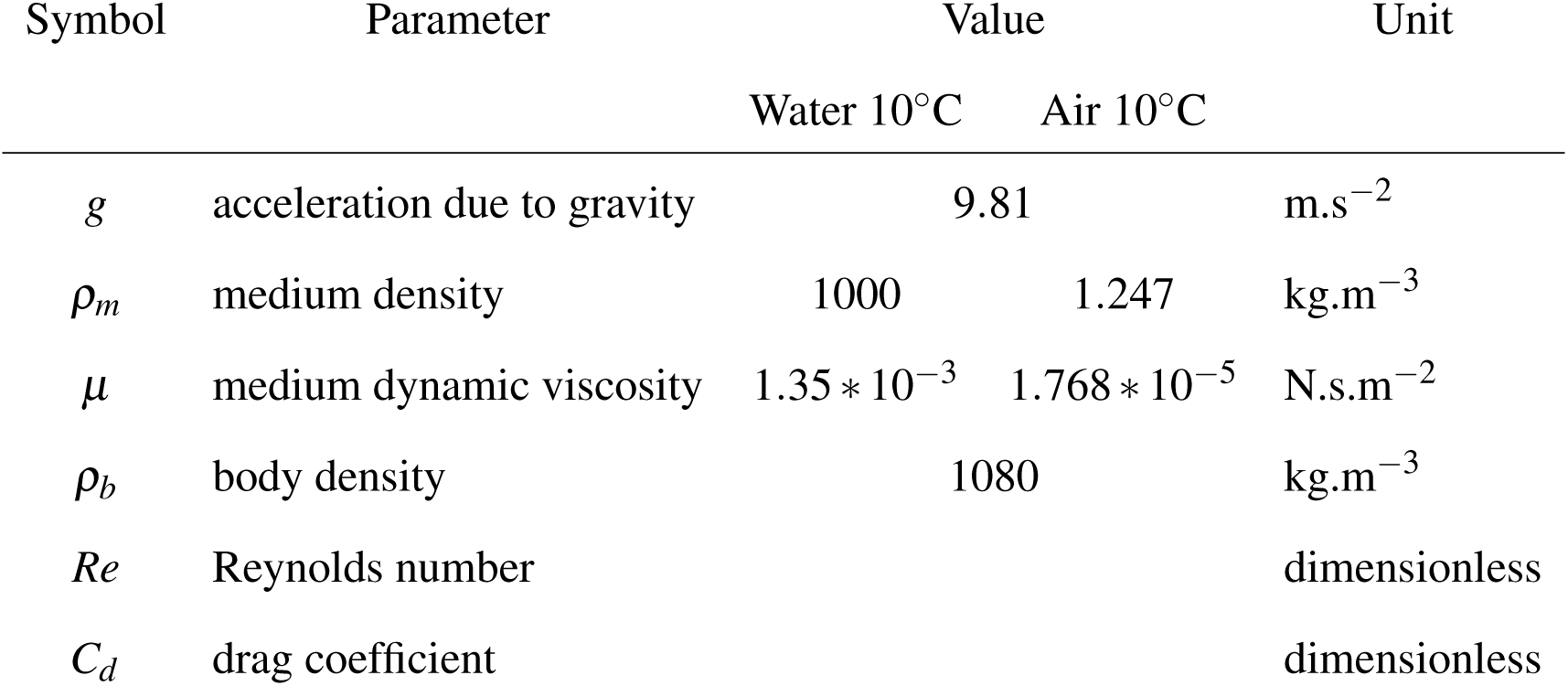
Physical parameters.

### 2.2 Biological parameters used by the model

These parameters are estimated using well-known allometric relationships (see Supplementary Table S1). Real data points have been used to calibrate some parameters. These data points are different from those that were used to test the model predictions.

#### 2.2.1 Energetic content

If the predator is able to find, capture and consume the prey, this predator will receive energy, which depends on the prey ash-free dry mass:

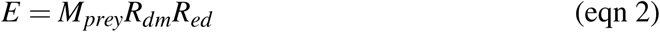

Where *R_dm_* is the ash-free dry mass to wet mass ratio, set as 0.16 (Ricciardi & Bourget, 1998), and *R_ed_* is energy to ash-free dry mass ratio, set as 23 ∗ 10^6^ J.kg^−1^ (Salonen, Sarvala, Hakala, & Viljanen, 1976).

#### 2.2.2 Metabolic rate

Each predator has a metabolic expenditure *per time* (*C_met_*) that scales with body mass. To allow for energetic expenditure due to muscular effort, the field metabolic rate is used (Savage et al., 2004; Hudson, Isaac, & Reuman, 2013):

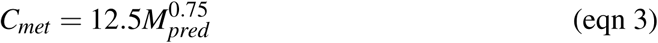

Parameters were estimated from data (Hudson, Isaac, & Reuman, 2013).

#### 2.2.3 Maximal muscular output and stroke period

The maximal muscular output (*F_Max_*) that a predator can develop scales with its body mass (Marden & Allen, 2002):

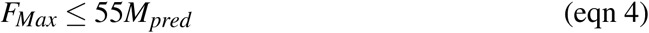

The time during which muscular forces are applied during motion, the stroke period (*t _f orce_*), scales with body size:

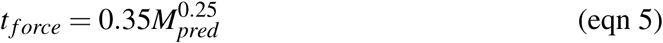

This parameter was estimated from real observations of species-specific speeds (Dodson, Ryan, Tollrian, & Lampert, 1997; Leis & Carson-Ewart, 1997; McDonald & Grünbaum, 2010). Equations 4 and 5 are similar for a prey.

#### 2.2.4 Detection distance

A predator can detect a prey individual (and the prey can detect the predator) within its detection distance. A larger species should have a larger detection sphere. We used a published model (Pawar, Dell, & Savage, 2012)

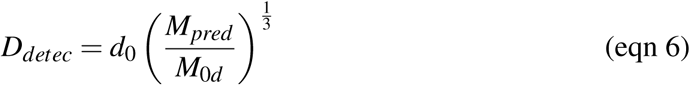

where *d*_0_ is the detection distance at reference size (set at 0.225 m), *M*_0*d*_ is the reference mass (set at 0.0376 kg). *d*_0_ and *M*_0_ were estimated by regression from Pawar, Dell, & Savage (2012). Prey detection distance was calculated in the same way.

### 2.3 Framework for calculation of speed and work

Predation is broken down into three different processes (search, capture, handling) involving motion, which lead to three different costs (one for each process). Calculations of these costs are all based on the same framework, where speed and cost are estimated using classical laws of Newtonian mechanics and fluid dynamics. However, the model assumes that species optimize different parameters for each predation process.

#### 2.3.1 Rationale for calculation

Although animal motion is diverse, it can be represented as an oscillatory movement (Bejan & Marden, 2006), a pattern observed in swimming, running or flying animals. Following this idea, we define a general framework for species motion.

Considering one oscillation, motion can be decomposed into a vertical and a horizontal component (Figure 2). Both are essential. The horizontal component represents the distance traveled between two points. However, this horizontal motion requires a vertical motion that either lifts the body or the surrounding medium (Bejan & Marden, 2006). The muscular output creates a force that is split between these two components.

**Figure 2:**
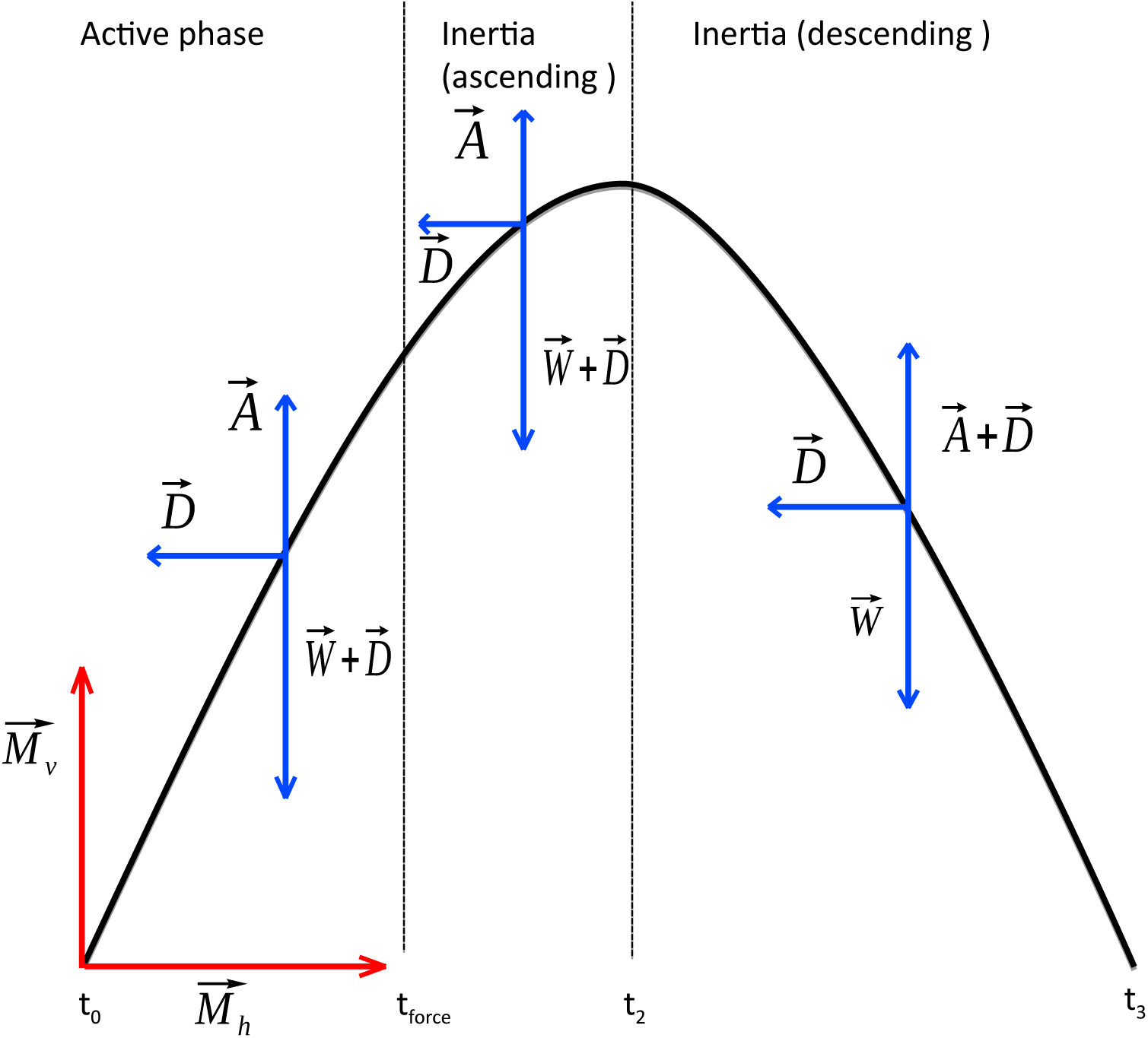
Framework for the calculation of motion cost. Motion is represented as an oscillation. An individual’s body moves upwards, then downwards, while moving forward. Red arrows represent mechanical forces applied by the organism during the stroke period (from *t*_0_ to *t _f orce_*). Blue arrows are external forces due to the surrounding medium: Archimedes’ force 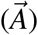, weight 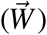, and drag 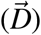. Direction of arrows account for the component of motion they affect (horizontal or vertical component). A given oscillation is split into three phases: an active phase, where a mechanical force is applied by the body, then an inertial phase, where the body pursues its motion upwards until its stops, and last an inertial (descending) phase, where the body returns to its original vertical position.

The vertical motion sequence can be decomposed into three successive phases. The first one is the active phase during which a muscular force (*F_Mv_*) is applied during the stoke period (see equations 4 and 5). The body is lifted by this muscular force (and Archimedes’ force) against its weight (due to gravity) and drag. The second phase is an inertial ascending phase: the organism pursues its lift by inertia until it stops. The last phase (inertial descending phase) occurs when the organism falls (or sinks) passively back to its original vertical position.

The horizontal motion sequence includes two phases. The first one is the active phase during which a muscular force (*F_Mh_*) allows a displacement of the body. This force is applied during the stoke period. The second phase is an inertial phase: the organisms pursues its motion until it stops.

The total time (*t*_3_, Figure 2) for both vertical and horizontal components is the same, as well as the stroke period (*t _f orce_*). It explains why the allocation of muscular force between the two components has a strong impact because a total allocation to the vertical component is useless, since the body stays at the same place horizontally, while a total allocation to the horizontal component is inefficient, since the organism cannot displace itself or the medium to move forward.

#### 2.3.2 Force allocation and work

For each of the predatory sequences (search, capture, handling), the aforementioned framework allows the calculation of the muscular force spent by the organism as well as the distance covered (see Supplementary Methods 2 for full details). Then, knowing the forces (*F_Mv_* + *F_Mh_*) and the distance covered during the active phase in both the vertical (*x_v_*) and horizontal (*x_h_*) plans, a work can be calculated, which is the energetic cost of motion.

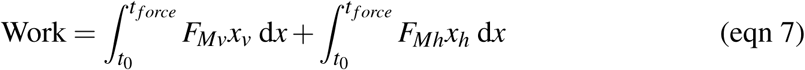

This work can be divided by the time of a whole oscillation (from *t*_0_ to *t*_3_) to yield a cost per unit time (Cost_*pt*_).

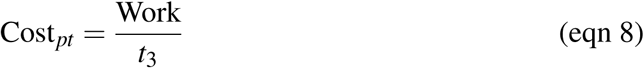

### 2.4 Calculation of search, capture and handling costs

For each predation cost (i.e., searching, capture and handling costs), force allocation between the vertical and horizontal components are estimated using an optimization procedure based on the Simplex method (Nelder & Mead, 1965).

#### 2.4.1 Searching cost

Searching cost represents energy spent by a predator to find its prey. It is based on a species-specific speed (*v̅*), which is the average speed throughout a whole oscillation. This speed is assumed to be sustainable for a long period of time. Thus, it optimizes the horizontal distance traveled for a minimal cost.

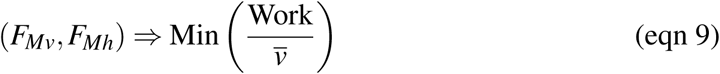

The instantaneous speed is greater when the muscular force is applied, and then decreases. Thus, an average speed gives a fair estimate of a cyclic process.

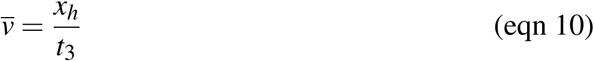

The optimization yields a species-specific speed that increases with body size.

To be consistent throughout the study, prey is assumed to fill 1% of the total volume of the medium (White, Ernest, Kerkhoff, & Enquist, 2007). Therefore, small prey are more abundant than large prey. Searching time is assumed to be the inverse of encounter rate (*E_r_*, see Supplementary Methods 3). Searching cost is the sum of the mechanical and metabolic costs (see above) during the search of a prey.

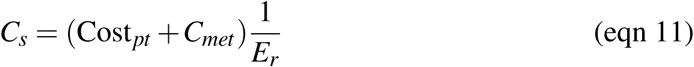

#### 2.4.2 Capture cost

To keep it as simple as possible, a capture sequence is based on a unique oscillation: the predator jumps and tries to seize the prey. The prey jumps and tries to escape the predator. This assumption is based on the observation that many predators do not actually pursue their prey during a long period of time; predators usually try to capture the prey quickly, and stop if they fail (Weihs & Webb, 1984).

The predator tries to optimize the horizontal distance (*x_h_*) covered during a unique jump.

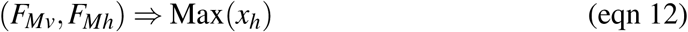

The predator may fail to capture the prey. Hence, a capture probability (*P_suc_*) is calculated. The prey can detect the predator if it is closer than the prey detection distance *D_prey_*, which is assumed to be the distance between the predator and its prey when the sequence starts.

First, the predator must cover the distance (*D_prey_*) between itself and its prey before it stops, otherwise the probability of capture is 0 (*P_suc_* = 0). Second, the relative speed between the predator (*v_Pred_*) and the prey (*v_Prey_*) at contact plays an essential role because if the prey is not able to move anymore, while the predator can pursue its motion, the probability of capture should be high. On the other hand, if the predator is at the end of its jump, while the prey can pursue its motion, the probability of capture should be low. We use a logistic function to describe this process:

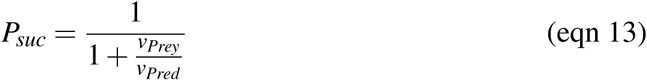

We assume that if *v_Pred_* = 0, it means that the predator is unable to cover the distance (*P_suc_* = 0).

The capture cost is paid by the predator no matter whether capture is successful or not. The number of attempts before a success is assumed to be the inverse of capture probability. The metabolic expenditure is paid for the duration of each jump (*t_c_*). Thus, the capture cost to effectively capture one prey is

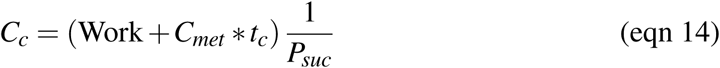

If *P_suc_* = 0, this predator-prey interaction is not feasible.

#### 2.4.3 Handling cost

The mechanical handling cost is based on the idea that a predator living in the water or air column has to maintain prey body mass during handling, otherwise it would lose its prey. Handling time depends on both predator and prey sizes (see Supplementary Methods 4).

Using the framework explained above, the predator body moves downwards due to gravity, and energy is spent periodically to lift its body to its original vertical position. The predator supports both its own mass and that of the prey. Only the vertical component of motion is used. Therefore, the cost *per time* (equation 8) becomes

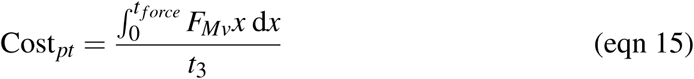

Handling cost is the sum of muscular and metabolic energy expenditure during handling time (*t_h_*):

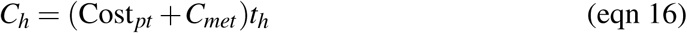

If the predator cannot lift its body to its original vertical position while carrying the prey, the interaction is assumed to be not feasible.

### 2.5 Size-related foraging costs and foraging limits

Each foraging cost (for searching, capturing, and handling the prey) varies with predator and prey sizes (Supplementary Figure S2). Each cost constrains the range of prey that a predator can consume, defining foraging limits. These limits can be either energetic or mechanical.

#### 2.5.1 Energetic limits

Energetic limits occur when a prey does not provide enough energy compared with the costs associated with its consumption. Limits are calculated for each foraging cost separately.

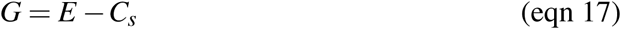

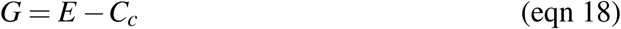

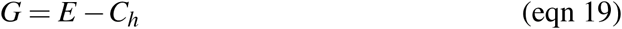

There is a limit for search (equation 17), capture (equation 18) and handling (equation 19). A specific energetic limit can be defined by assuming that metabolism is the only cost.

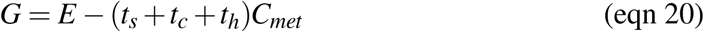

For each predator and for each cost, the smaller prey leading to *G* > 0 defines the corresponding energetic limit for this cost, which represents the minimal prey size that allows a given predator to fulfill its energetic expenditure for predation, when only the corresponding cost is considered.

#### 2.5.2 Mechanical limits

Mechanical limits are due to a lack of sufficient muscular power for the predator. The capture mechanical limit occurs when the predator is unable to reach the prey irrespective of the number of attempts (i.e., *P_suc_* = 0). Similarly, the handling mechanical limit occurs when the predator is unable to lift the prey during handling time. In both cases, the predator-prey interaction is assumed not to be feasible, and a gain is impossible to calculate.

### 2.6 Use of empirical data

We compared model predictions with empirical data on predator-prey relationships (see Supplementary Methods 5). Data come from the most extensive database of predator and prey body sizes currently published (Brose et al., 2005), supplemented with data for flying predators that we collected directly from published articles. For our analysis and presentation in graphs, data points were sorted and grouped according to whether or not interactions fitted the model’s assumptions. These assumptions were that 1) the interaction occurs on a one-to-one base, 2) the predator tries to actively seize the prey, and the prey actively tries to escape the predator, and 3) both predator and prey can detect each other without interference (i.e., the predator cannot hide itself). Data that did not fit the model’s assumptions were sorted according to which assumption was violated or relaxed. If several assumptions were violated, we considered the most limiting one.

### 2.7 Model accuracy

In order to evaluate to what extent the model was able to predict feasible and non-feasible interactions, the True Skill Statistics (TSS) was computed (Allouche, Tsoar, & Kadmon, 2006) on some of the data points that fitted the model assumptions. For each food web, four metrics were calculated: *a*, the number of links predicted and observed, *b*, the number of links predicted with no corresponding observation, *c*, the number of links observed but predicted absent by the model, and *d*, the number of predicted and observed absence of links. TSS quantifies the amount of successful predictions relative to false predictions.

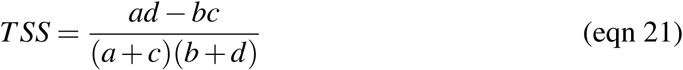

Its value ranges between 1 (perfect prediction) and −1 (inverted prediction).

We selected data that came from food web studies, and excluded some data that came from studies focusing on single predators.

## 3 Results

### 3.1 Predicted interactions

Combining a mechanical model of predation with metabolic laws allowed us to calculate the net energy gains of a predator consuming a prey item of a given body size. Three cases can occur: (1) if an interaction leads to a positive net energetic gain, it is considered feasible and sustainable; (2) if the interaction leads to a negative net energetic gain, it is considered feasible but unsustainable; (3) if the predator cannot capture the prey, the interaction is considered unfeasible.

We found that each predator can feed on a range of prey sizes that varies with its body size. Typically, larger predators feed on larger prey, as is often observed in nature (Figure 3). The model predicts that predators should be larger than their prey, and this constraint is stronger for flying than pelagic predators. The gains of predators of similar sizes are also consistently lower in flying predators than in pelagic ones. The prey giving the highest net energetic gain is always the largest prey that a predator can consume.

**Figure 3:**
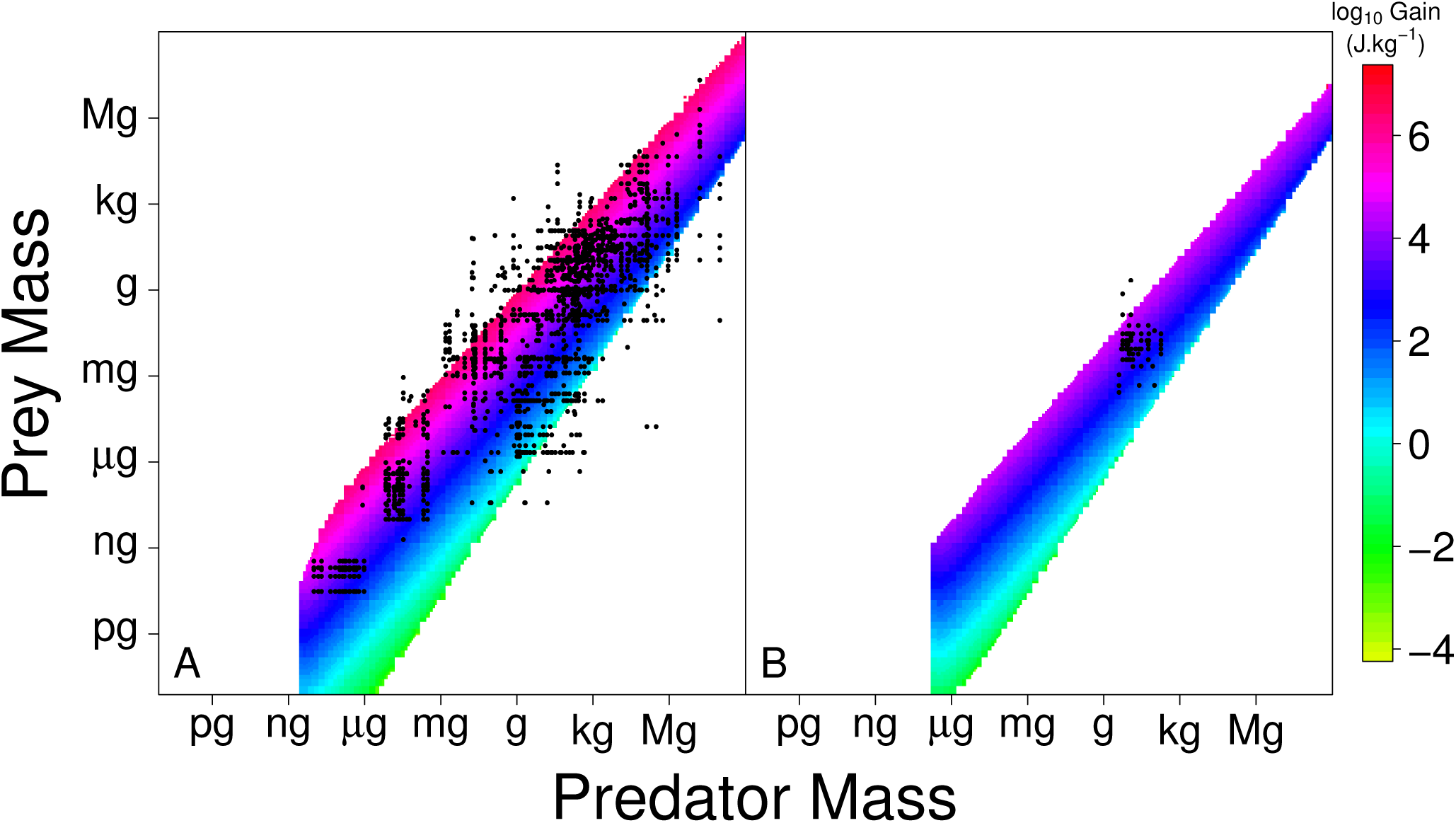
Net gains on predation for pelagic (A) and flying (B) predators. Heat maps show net energetic gains for predation on a one-to-one based interaction. Gains are weighed by predator mass in order to allow comparisons between predators. When predator size increases, prey size should also increase because larger predators can capture larger prey. However, small prey do not provide enough energy, and therefore they become not sustainable for large predators. Points represent real interactions that fit the model assumptions within different aquatic systems either in marine or freshwater habitats, and for flying predators. In aquatic systems predator size of the empirical data ranges from rotifers to whales; 80% of the points fall within the predicted range of prey sizes. Freshwater and salt water did not show any significant difference. Thus, these ecosystems are shown together. In air, data are restricted to insectivorous bats and birds since many flying predators come back on the ground during handling time; 96% of the points fall within the predicted range of prey size.

Despite its simplifying assumptions, the model predicts most observed interactions across the whole range of predator sizes (from zooplankton to large vertebrates). About 80% and 97% of pelagic and flying predators respectively, fall within the predicted range of values (Figure 3). Moreover, TSS shows that the model is able to predict a large part of the trophic links within food webs (Figure 4). The accuracy of the model was compared over the same data points with the parameterized niche model, which represents the most advanced attempt at predicting pairwise interactions based on an empirical approach that is using body size as the only matching trait (Gravel, Poisot, Albouy, Velez, & Mouillot, 2013). Our model often shows greater accuracy than the niche model.

**Figure 4:**
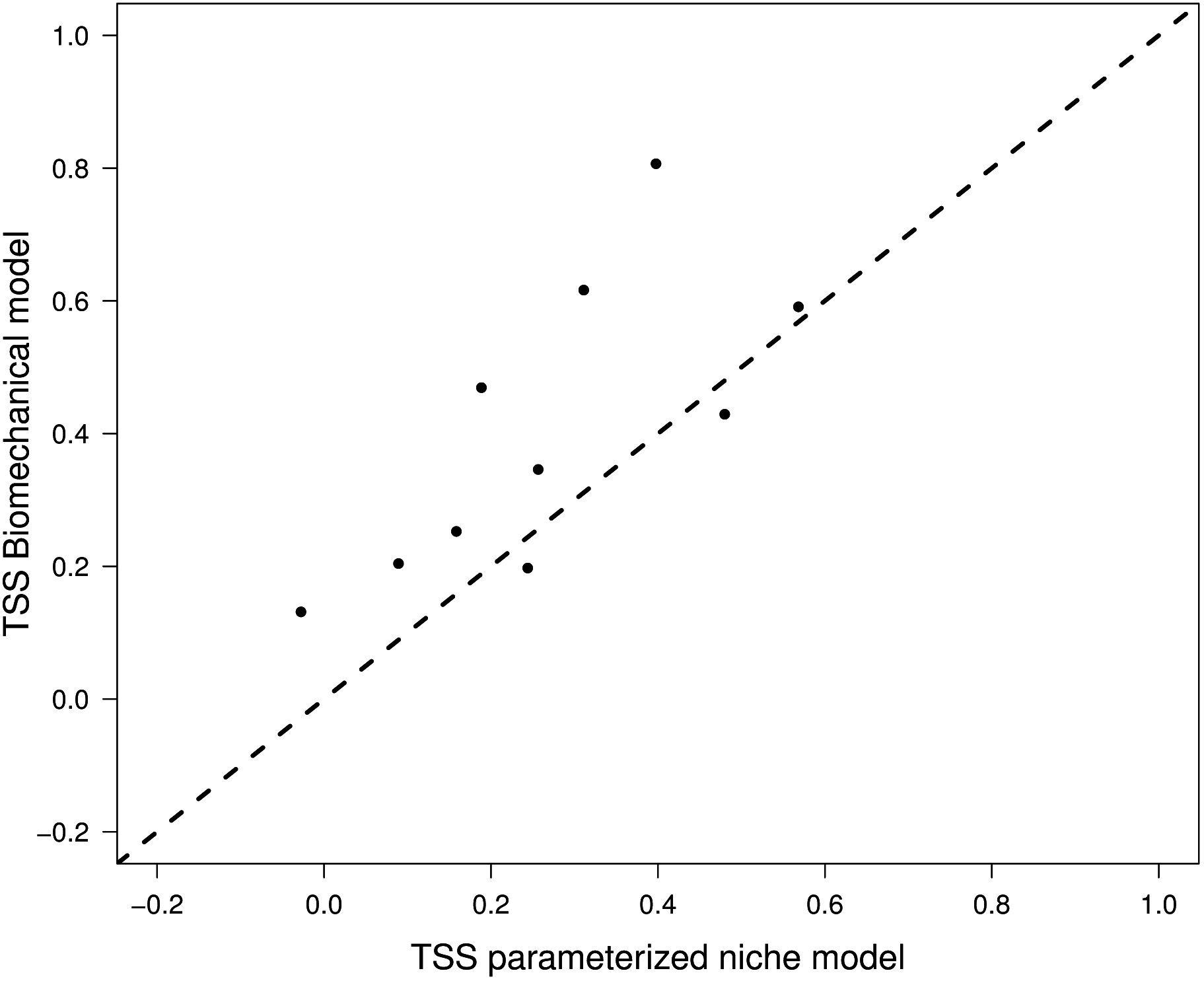
Comparison of True Skill Statistics (TSS) between our biomechanical model and a parameterized niche model on ten food webs. Each point is a food web; the dashed line is a 1:1 line (i.e., similar accuracy). Most points are located above the line, which means that the biomechanical model shows greater accuracy.

### 3.2 Foraging limits

In a second step, we analyzed our model in detail to determine how the various mechanical and energetic components of the model constrain the size of prey that a predator can consume. The maximum prey size that a predator can eat is determined by mechanical constraints. In fact, larger prey individuals can both detect a predator earlier and develop greater velocities (see Methods), resulting in successful escape. Thus, there is a maximum size for the prey that a predator of a given size can capture (solid blue lines on Figure 5). Another mechanical constraint is related to handling, when the prey is too large and the predator is unable to develop sufficient mechanical power to hover while maintaining its prey (solid red lines on Figure 5). With the set of parameter values we chose, which are typical of generic pelagic and airborne food webs (Table 1 and Supplementary Table S1), it is capture that mechanically constrains the upper prey size (Figure 5).

**Figure 5:**
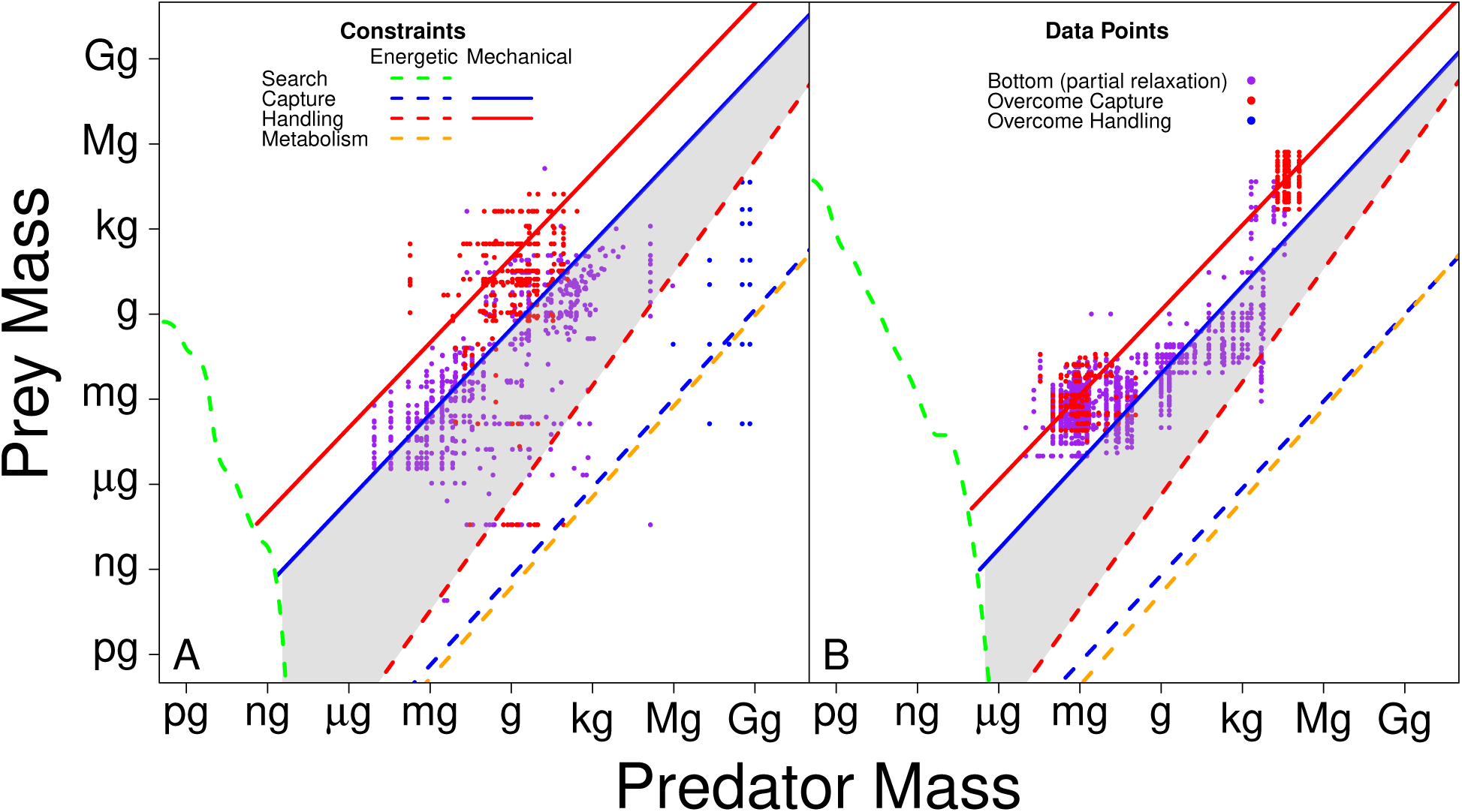
Constraints on feeding interactions in aquatic (A) and terrestrial (B) systems. Dashed lines are the energetic constraints (lower prey size allowing a positive net gain when the corresponding cost is the only one acting). Solid lines are the mechanical constraints (upper prey size that a predator can capture or handle under the model assumptions). Grey areas show the predicted interactions (Figure 3). Upper prey size is determined by capture mechanical constraint. Lower prey size is mostly constrained by handling energetic constraint. Lower predator size is mostly determined by searching cost. Colour of data points shows which constraint is relaxed. Red points are predators that overcome the mechanical capture constraint. Blue points are predators that overcome handling energetic constraint by consuming several small prey at a time. Purple points are predators living on a hard surface (bottom of aquatic systems or ground). They relax capture mechanical constraint since they can hide themselves, and they relax handling energetic constraint since they do not need to carry the prey. These points include flying predators that return to the ground to handle their prey.

In contrast, minimum prey size is limited by net energy gain. The amount of energy given by a prey increases with its size (Supplementary Figure S2). Hence, small prey sizes are poor energetic rewards for predators. Searching, capture, handling, and metabolic costs may further decrease prey profitability. Handling cost (dashed red lines on Figure 5) is the most limiting cost for predators larger than a few *μ*g because the cost of hovering dominates over the other costs that are capture cost (dashed blue lines) and metabolic cost (dashed orange lines). For pelagic predators that are below the nanogram range (below the ?g range for flying predators), searching cost is the limiting cost (green dashed lines on Figure 5). Small predators have short detection distances and low velocities resulting in too rare encounters under the prey densities assumed in the model (see Methods).

### 3.3 Overcoming foraging limits

Not all predators in our dataset pay the full costs of searching, capturing and handling their prey. Some predators overcome the capture mechanical limit (red points on Figure 5) by feeding on prey that do not move (e.g., on sponges or corals) or that move at a lower speed than expected according to their size (e.g., on gastropods). Such predators should be limited in their choice of prey by handling, the next process to act on the range of feasible prey sizes.

Other predators decrease the energetic cost of handling, which is mainly the cost of hovering in the case of small prey, by consuming several small prey items at a time, such as strikingly performed by plankton-feeding whales (blue points on Figure 5). Finally, some predators overcome both the capture and handling limitations by living on the bottom (benthic, running or crawling predators; purple points on Figure 5). Such predators spend less energy in managing their buoyancy while handling their prey. Many flying predators, insects and birds, move to a hard surface during the handling of their prey, and thus belong to this category of predators. Since surfaces generally bear complex landscapes, predators can hide and come closer to their prey before being detected, which increases the likelihood of capture. Such predators have potentially no limits to the maximum size for the prey they can capture, in particular if they hunt in group. Our results however, show that this category of predators target prey with maximum prey sizes that are very close to the handling mechanical constraint (solid red lines on Figure 5).

## 4 Discussion

This study presents a mechanistic and mechanical model that predicts the occurrence of an interaction between a predator and a prey species with specified body sizes. For each predator size, we calculate the feasibility and energetic profitability from eating a prey of a given size, using a Newtonian, mechanical model associated with general metabolic laws. The predicted size ranges of feasible and profitable predator interactions compare well with observed interactions, as documented in the most extensive size-based predation database published so far (Brose et al., 2005), augmented with additional data on flying predators. Since our model is mechanistic and is based on general laws, it neither needs calibration for each food web, nor any extra-information (such as a co-occurrence matrix). Nonetheless, the model shows a greater accuracy than a parameterized niche model (Gravel, Poisot, Albouy, Velez, & Mouillot, 2013). Both models usually overestimate the number of observed links, which is quite a common issue. But it is difficult to sample all links of a given food web (Martinez, Hawkins, Dawah, & Feifarek, 1999), therefore an absence of observation does not necessarily mean an absence of link.

There are a number of other mechanistic models of food webs (Loeuille & Loreau, 2005; Petchey, Beckerman, Riede, & Warren, 2008; Carbone, Codron, Scofield, Clauss, & Bielby, 2014). Among these models, the allometric diet breadth model (ADBM) is the only model that, like ours, aims at predicting realized predation interactions, rather than at simulating virtual food webs (Petchey, Beckerman, Riede, & Warren, 2008). ADBM adopts an approach that is similar to ours in many respects. Both models predict the diet of individual predator species based on body size as the main trait, and on a mechanistic model describing the energy gain from the prey. The choice of the mechanistic underpinnings is where the two models diverge: we base our calculations on a combination of mechanical and metabolic laws; ADBM is based on optimal foraging theory (Beckerman, Petchey, & Warren, 2006). Rather than confront the two models, we see them as complementary. ADBM uses empirical allometric relationships, whose parameters need to be estimated from the food web datasets examined, to include body size as a trait in the model; we offer a mechanistic derivation of these allometries. ADBM does not subtract energetic costs from the energy content of the prey; we account for the costs related to the search, capture and handling of the prey. On the other hand, our model does not offer a ranking in the choice of prey, only net gain estimates; ADBM offers a ranking of species based on optimal foraging. Thus, we see the next obvious step in the development of our model in the combination of the two modeling approaches.

Our model matches some of the common body size patterns observed in food webs (Tucker & Rogers, 2014). In particular, predators consume smaller prey in air than in water, but the patterns remain similar otherwise. Thus, constraints due to mechanical factors are stronger in air, but apply in the same way as in water. There is also a greater number of predators that handle their prey in the water column, compared with the number of flying predators that handle their prey in the air (compare numbers of data points between the 2 panels of Figure 3). Our model provides an explanation for this difference: hovering costs are lower in the water column than in the air, due to higher buoyancy. Moreover, the bottom is generally farther from the vertical position of pelagic predators (in oceans and large lakes), requiring a significant energy expenditure to be reached. In contrast, it is easier for flying predators to return to the ground during handling, to a degree that we could only find a few insectivorous bats and birds, as well as bat hawks, that matched the assumption of continuous hovering in the air (Figure 3b). More generally, since hovering is easier in water than in air, predator motion during capture has wider amplitude in water, which leads to a greater chance for the predator to reach its prey. It explains why, in air, predators are more constrained by the capture mechanical limit than are aquatic predators.

Tucker & Rogers (2014) found that predator-prey body size ratios are generally greater for carnivores than for herbivores. Usually, herbivores consume resources that do not move or move slowly compared with their size, so that they are able to overcome the mechanical constraints set by the capture process. Carnivores face stronger mechanical capture limit because the prey can escape. Thus, a carnivore has better chances to capture a small prey than a large one, which leads to a larger body size ratio. In summary, our model offers a unique opportunity for a unified understanding of predator-prey patterns across habitats and trophic levels.

Despite the overall good performance of the model, we see that predators often prey on organisms that the model considers smaller than the optimal size. We think that this mainly results from our use of generic, simplified allometric equations to describe important parameters in the model, such as prey population densities, maximum accelerations, and detection distances. Recent advances in the field of allometry have shown that the effect of body size can be more complicated than previously acknowledged (Pawar, Dell, & Savage, 2012; Wilson et al., 2015; Hirt, Jetz, Rall, & Brose, 2017), although it is still predictable (Kiørboe & Hirst, 2014). Our model’s predictability would certainly benefit from an increase in the realism of the allometric equations it uses.

There are other factors that may lead to a suboptimal choice of prey in real ecosystems. The optimal prey might be absent, or show defense traits that make it challenging to find, capture or handle. Several studies have shown that further functional traits besides body size are necessary for an accurate prediction of trophic interactions (Eklöf et al., 2013; Blanchard, Heneghan, Everett, Trebilco, & Richardson, 2017). However, which traits need to be included first is yet debated (Boukal, 2014). Based on our model, and in agreement with other authors (Higham, 2007; Boukal, 2014), we propose as likely candidates the biomechanical traits related to predator and prey performances, after accounting for the effect of body size, i.e., deviations from allometries in velocities, accelerations and muscular forces.

There are also predators that feed on prey with body sizes beyond the predicted range of prey sizes. Such predators probably evolved strategies to get past the capture and handling mechanical limits. One important strategy is the ambush or sit-and-wait strategy (Kiørboe, 2011), which leads to the capture of larger prey than expected in terrestrial ecosystems (Supplementary Figure S3). Our model suggests that the largest prey size for these predators is set by the handling mechanical limit (but not for web-weaving spiders). Predators differ in other aspects of their search (Bläßle & Tyson, 2016), capture (Higham, 2007) and handling strategies (Kiørboe, 2011). Building mechanical models with a similar approach to ours for most of the major predation strategies would certainly advance our understanding of food web structure and its predictability.

Our model describes the interaction between a predator of a given size and a prey of a given size at a given moment in time, and it looks at the energetic balance between costs and gains during the predation act. But a predator usually needs to share its time between predation and other activities such as reproduction, recovery and the avoidance of its own predators. The energy gained from the predation act must also be spent in these activities. Our model ignores these additional energetic costs for the time being. The minimum prey size resulting in a positive net energy gain should be higher when all activities of the predator are included. It is far from obvious to calculate the energy cost related to the various activities of a predator. However, some existing allometric works open the door to such a development in our model (Preisser & Orrock, 2012; Rizzuto, Carbone, & Pawar, 2017).

Despite the high level of abstraction of our model, it fits empirical data remarkably well. This suggests that predator-prey interactions in pelagic and aerial habitats are heavily constrained by mechanical factors despite hundreds of millions of years of evolution. It seems that numerous species follow the assumptions of the model and stay within the limits imposed by mechanical and energetic constraints, while other species have adapted to overcome these limits in a way that is consistent with our model, albeit with relaxed assumptions (Figure 5). Overall, this suggests that physical factors have played a major role in the evolution of trophic interactions. Our model offers a general framework for the study of the mechanical bases of trophic interactions across a wide range of body sizes. It also provides general conclusions and mechanisms underpinning well-known empirical patterns in the structure of food webs beyond apparent discrepancies between media. Our work strongly emphasizes the need to consider the physical medium to understand the ecology of food webs (Denny, 2016). In that sense, it is an ecosystem approach at heart, one that does not separate the organisms “from their special environments, with which they form one physical system” (Tansley, 1935).

## Acknowledgements

Authors thank Dr. U. Brose and Dr. D. Gravel for their useful comments on an earlier version of the paper. ML was supported by the TULIP Laboratory of Excellence (ANR-10-LABX-41). GFF was supported by an NSERC Discovery Grant. MC and SP thank the Gustaf & Hanna Winblad fund for the advancement of zoological research.

## Data accessibility

Data used in the present manuscript will be archived in a public repository (Dryad).

The R code used to implement the model and test its accuracy will be archived in a public repository (Zenodo).

## References

Allesina, S. & Tang, S. (2012). Stability criteria for complex ecosystems. Nature, 483(7388), 205–208. doi:10.1038/nature10832.

Allouche, O., Tsoar, A., & Kadmon, R. (2006). Assessing the accuracy of species distribution models: prevalence, kappa and the true skill statistic (TSS). Journal of Applied Ecology, 43(6), 1223–1232. doi:10.1111/j.1365-2664.2006.01214.x.

Beckerman, A. P., Petchey, O. L., & Warren, P. H. (2006). Foraging biology predicts food web complexity. Proceedings of the National academy of Sciences of the United States of America, 103(37), 13745–13749. doi:10.1073/pnas.0603039103.

Bejan, A. & Marden, J. H. (2006). Unifying constructal theory for scale effects in running, swimming and flying. Journal of Experimental Biology, 209(Pt 2), 238–248. doi:10.1242/jeb.01974.

Blanchard, J. L., Heneghan, R. F., Everett, J. D., Trebilco, R., & Richardson, A. J. (2017). From Bacteria to Whales: Using Functional Size Spectra to Model Marine Ecosystems. Trends in ecology and evolution, 32(3), 174–186. doi:10.1016/j.tree.2016.12.003.

Bläßle, A. & Tyson, R. (2016). First capture success in two dimensions: The search for prey by a random walk predator in a comprehensive space of random walks. Ecological Complexity, 28, 24–35. doi:10.1016/j.ecocom.2016.09.004.

Boukal, D. S. (2014). Trait- and size-based descriptions of trophic links in freshwater food webs: Current status and perspectives. Journal of Limnology, 73(1 SUPPL), 171–185.

Brose, U. (2010). Body-mass constraints on foraging behaviour determine population and food-web dynamics. Functional Ecology, 24(1), 28–34. doi:10.1111/j.1365-2435.2009.01618.x.

Brose, U., Cushing, L., Berlow, E. L., Jonsson, T., Banasek-Richter, C., Bersier, L.-F., Blan-chard, J. L., Brey, T., Carpenter, S. R., Blandenier, M.-F. C., Cohen, J. E., Dawah, H. A., Dell, T., Edwards, F., Harper-Smith, S., Jacob, U., Knapp, R. A., Ledger, M. E., Memmott, J., Mintenbeck, K., Pinnegar, J. K., Rall, B. C., Rayner, T., Ruess, L., Ulrich, W., Warren, P., Williams, R. J., Woodward, G., Yodzis, P., & Martinez, N. D. (2005). Body Sizes Of Consumers And Their Resources. Ecology, 86(9), 2545–2545. doi:10.1890/05-0379.

Carbone, C., Codron, D., Scofield, C., Clauss, M., & Bielby, J. (2014). Geometric factors influencing the diet of vertebrate predators in marine and terrestrial environments. Ecology Letters, 17(12), 1553–1559. doi:10.1111/ele.12375.

Cohen, J. E. & Newman, C. M. (1985). A Stochastic Theory of Community Food Webs: I. Models and Aggregated Data. Proceedings of the Royal Society B: Biological Sciences, 224(1237), 421–448. doi:10.1098/rspb.1985.0042.

Cohen, J. E., Pimm, S. L., Yodzis, P., & Saldaña, J. (1993). Body Sizes of Animal Predators and Animal Prey in Food Webs. Journal of Animal Ecology, 62(1), 67–78.

Denny, M. W. (2016). Ecological Mechanics: Principles of Life’s Physical Interactions. Princeton University Press.

Dodson, S. I., Ryan, S., Tollrian, R., & Lampert, W. (1997). Individual swimming behavior of *Daphnia*: effects of food, light and container size in four clones. Journal of Plankton Research, 19(10), 1537–1552. doi:10.1093/plankt/19.10.1537.

Eklöf, A., Jacob, U., Kopp, J., Bosch, J., Castro-Urgal, R., Chacoff, N. P., Dalsgaard, B., de Sassi, C., Galetti, M., Guimarães, P. R., Lomáscolo, S. B., Martín González, A. M., Pizo, M. A., Rader, R., Rodrigo, A., Tylianakis, J. M., Vázquez, D. P., & Allesina, S. (2013). The dimensionality of ecological networks. Ecology Letters, 16(5), 577–583. doi:10.1111/ele.12081.

Evans, D. M., Kitson, J. J. N., Lunt, D. H., Straw, N. A., & Pocock, M. J. O. (2016). Merging DNA metabarcoding and ecological network analysis to understand and build resilient terrestrial ecosystems. Functional Ecology, 30(12), 1904–1916. doi:10.1111/1365-2435.12659.

Gravel, D., Poisot, T., Albouy, C., Velez, L., & Mouillot, D. (2013). Inferring food web structure from predator-prey body size relationships. Methods in Ecology and Evolution, 4(11), 1083–1090. doi:10.1111/2041-210X.12103.

Higham, T. E. (2007). The integration of locomotion and prey capture in vertebrates: Morphology, behavior, and performance. Integrative and Comparative Biology, 47(1), 82–95. doi:10.1093/icb/icm021.

Hirt, M. R., Jetz, W., Rall, B. C., & Brose, U. (2017). A general scaling law reveals why the largest animals are not the fastest. Nature Ecology & Evolution, 1(8), 1116.

Hudson, L. N., Isaac, N. J. B., & Reuman, D. C. (2013). The relationship between body mass and field metabolic rate among individual birds and mammals. Journal of Animal Ecology, 82(5), 1009–1020. doi:10.1111/1365-2656.12086.

Kiørboe, T. (2011). How zooplankton feed: mechanisms, traits and trade-offs. Biological Reviews, 86(2), 311–339. doi:10.1111/j.1469-185X.2010.00148.x.

Kiørboe, T. & Hirst, A. G. (2014). Shifts in mass scaling of respiration, feeding, and growth rates across life-form transitions in marine pelagic organisms. The American naturalist, 183(4). doi:10.1086/675241.

Leis, J. & Carson-Ewart, B. (1997). In situ swimming speeds of the late pelagic larvae of some Indo-Pacific coral-reef fishes. Marine Ecology Progress Series, 159, 165–174. doi:10.3354/meps159165.

Loeuille, N. & Loreau, M. (2005). Evolutionary emergence of size-structured food webs. Proceedings of the National academy of Sciences of the United States of America, 102(16), 5761–5766. doi:10.1073/pnas.0408424102.

Marden, J. H. & Allen, L. R. (2002). Molecules, muscles, and machines: Universal performance characteristics of motors. Proceedings of the National academy of Sciences of the United States of America, 99(7), 4161–4166. doi:10.1073/pnas.022052899.

Martinez, N. D., Hawkins, B. A., Dawah, H. A., & Feifarek, B. P. (1999). Effects Of Sampling Effort On Characterization Of Food-Web Structure. Ecology, 80(3), 1044–1055. doi:10.1890/0012-9658(1999)080[1044:EOSEOC]2.0.CO;2.

McDonald, K. A. & Grünbaum, D. (2010). Swimming performance in early development and the “other” consequences of egg size for ciliated planktonic larvae. Integrative and comparative biology, 50(4), 589–605. doi:10.1093/icb/icq090.

Nelder, J. A. & Mead, R. (1965). A Simplex Method for Function Minimization. The Computer Journal, 7(4), 308–313. doi:10.1093/comjnl/7.4.308.

Pawar, S., Dell, A. I., & Savage, V. M. (2012). Dimensionality of consumer search space drives trophic interaction strengths. Nature, 486(7404), 485–489. doi:10.1038/nature11131.

Petchey, O. L., Beckerman, A. P., Riede, J. O., & Warren, P. H. (2008). Size, foraging, and food web structure. Proceedings of the National academy of Sciences of the United States of America, 105(11), 4191–4196. doi:10.1073/pnas.0710672105.

Preisser, E. L. & Orrock, J. L. (2012). The allometry of fear: interspecific relationships between body size and response to predation risk. Ecosphere, 3(9), art77. doi:10.1890/ES12-00084.1.

Ricciardi, A. & Bourget, E. (1998). Weight-to-weight conversion factors for marine benthic macroinvertebrates. Marine Ecology Progress Series, 245–251.

Rizzuto, M., Carbone, C., & Pawar, S. (2017). Foraging constraints reverse the scaling of activity time in carnivores. Nature Ecology & Evolution, 1. doi:10.1038/s41559-017-0386-1.

Romanuk, T. N., Zhou, Y., Brose, U., Berlow, E. L., Williams, R. J., & Martinez, N. D. (2009). Predicting invasion success in complex ecological networks. Philosophical Transactions of the Royal Society of London B: Biological Sciences, 364(1524), 1743–1754.

Salonen, K., Sarvala, J., Hakala, I., & Viljanen, M.-L. (1976). The relation of energy and organic carbon in aquatic invertebrates1. Limnology and Oceanography, 21(5), 724–730. doi:10.4319/lo.1976.21.5.0724.

Savage, V. M., Gillooly, J. F., Woodruff, W. H., West, G. B., Allen, A. P., Enquist, B. J., & Brown, J. H. (2004). The predominance of quarter-power scaling in biology. Functional Ecology, 18(2), 257–282. doi:10.1111/j.0269-8463.2004.00856.x.

Stouffer, D. B. (2010). Scaling from individuals to networks in food webs. Functional Ecology, 24(1), 44–51. doi:10.1111/j.1365-2435.2009.01644.x.

Tansley, A. G. (1935). The use and abuse of vegetational concepts and terms. Ecology, 16(3), 284–307.

Thompson, R. M., Brose, U., Dunne, J. A., Hall, R. O., Hladyz, S., Kitching, R. L., Martinez, N. D., Rantala, H., Romanuk, T. N., Stouffer, D. B., & Tylianakis, J. M. (2012). Food webs: reconciling the structure and function of biodiversity. Trends in ecology & evolution, 27(12), 689–697. doi:10.1016/j.tree.2012.08.005.

Tucker, M. A. & Rogers, T. L. (2014). Examining predator-prey body size, trophic level and body mass across marine and terrestrial mammals. Proceedings of the Royal Society B - Biological Sciences, 281, 20142103.

Webb, T. J. (2012). Marine and terrestrial ecology: unifying concepts, revealing differences. Trends in ecology and evolution, 27(10), 535–541.

Weihs, D. & Webb, P. (1984). Optimal avoidance and evasion tactics in predator-prey interactions. Journal of Theoretical Biology, 106(2), 189–206. doi:10.1016/0022-5193(84)90019-5.

White, E. P., Ernest, S. M., Kerkhoff, A. J., & Enquist, B. J. (2007). Relationships between body size and abundance in ecology. Trends in ecology and evolution, 22(6), 323–330. doi:10.1016/j.tree.2007.03.007.

Williams, R. J. & Martinez, N. D. (2000). Simple rules yield complex food webs. Nature, 404(6774), 180–183. doi:10.1038/35004572.

Williams, R. J. & Martinez, N. D. (2008). Success and its limits among structural models of complex food webs. Journal of Animal Ecology, 77(3), 512–519. doi:10.1111/j.1365-2656.2008.01362.x.

Wilson, R. P., Griffiths, I. W., Mills, M. G., Carbone, C., Wilson, J. W., & Scantlebury, D. M. (2015). Mass enhances speed but diminishes turn capacity in terrestrial pursuit predators. eLife, 4, e06487. doi:10.7554/eLife.06487.

Yvon-Durocher, G., Reiss, J., Blanchard, J., Ebenman, B., Perkins, D. M., Reuman, D. C., Thierry, A., Woodward, G., & Petchey, O. L. (2011). Across ecosystem comparisons of size structure: methods, approaches and prospects. Oikos, 120(4), 550–563. doi:10.1111/j.1600-0706.2010.18863.x.

